# Load-dependent amygdala neural representation and amygdala-hippocampus interaction in non-emotional working memory maintenance

**DOI:** 10.1101/2024.05.30.596374

**Authors:** Chenyang Li, Yulong Peng, Ruixue Wang, Xianhui He, Ying Cai, Yuehui Ma, Dengchang Wu, Minmin Wang, Shaomin Zhang

## Abstract

Retaining information in working memory (WM) is an active process that requires neural activity within and between regions. The human amygdala (AMY) and hippocampus (HPC) are known to play crucial roles in WM processing. Although electrophysiological studies have revealed that the HPC supports multi-item maintenance in a load-dependent manner, the characteristics of the AMY and the circuit-level mechanisms underlying AMY-HPC interactions remain largely unexplored. To address this knowledge gap, intracranial EEG recordings from the AMY and HPC in nine epileptic patients were employed to evaluate intraregional neural representations and interregional communications during maintenance under different non-emotional WM loads. High load enhanced low-frequency power and intraregional theta–gamma phase-amplitude coupling (PAC) in the AMY and HPC. At the network level, a high load elicited an increase in the strength of the HPC theta phase modulation, which entrains the AMY gamma amplitude. Furthermore, a high load increased AMY-anterior HPC (aHPC) theta phase synchrony and directional connectivity strength from the aHPC to the AMY. Conversely, posterior HPC (pHPC)-AMY synchrony was not affected by load variations. Overall, these findings highlight the importance of the AMY in non-emotional WM tasks and provide new insights into the neurophysiological basis of AMY-HPC interactions during WM maintenance.

## 1. Introduction

Working memory (WM), the ability to maintain and manipulate information over a short period,^1^ is crucial for higher-level cognitive functions.^2^ WM tasks comprise encoding, maintenance, and retrieval phases, with the maintenance phase serving as the central component and distinguishing WM from other types of memory.^3,4^ Maintenance is pivotal for successful WM performance because it requires maintaining active neural representations even in the absence of external stimuli.^5^ The amount of information to be retained, i.e., WM load, has been shown to affect neural processing.^6^ Exploring the effects of WM load can enhance our understanding of how the brain coordinates different regions to accomplish complex cognitive tasks. In humans, the amygdala (AMY) and hippocampus (HPC) are thought to be involved in WM processing.^7,8^ Although electrophysiological studies in humans have shown that the HPC supports non-emotional multi-item maintenance in a load-dependent manner, the role of the AMY and AMY-HPC interactions in supporting non-emotional WM maintenance and their cognitive load-dependent properties remain largely unexplored.

Recent studies have shown that the HPC is a critical region involved in WM maintenance, especially when handling an increasing cognitive load.^9–11^ While the AMY has traditionally been believed to be related primarily to emotions, recent findings indicate that it also participates in non-emotional WM. The multidimensional response characteristics of this region, such as blood oxygen level-dependent (BOLD) responses and sustained neural activity, are dependent on memory loads.^12,13^ In addition, activation of the AMY via electrical stimulation or peripheral drug administration could enhance non-emotional memory consolidation.^14–16^ These existing studies suggest that AMY activity is critical for modulating memory in the absence of emotional stimuli. Involvement of the AMY in WM maintenance was supported by evidence that the degree of phase locking in AMY-selective neurons is load dependent, with increased phase locking occurring under higher loads.^17^ However, another study failed to provide evidence supporting involvement of the AMY in maintenance, which was attributed to the inability of AMY neuronal activity to decode load levels during maintenance.^18^ Therefore, whether the AMY is involved in WM maintenance and shows load dependency is under intensive debate.

WM relies on coordinated processing between functionally interconnected brain regions.^19^ It has been extensively proposed that interactions between the AMY and HPC support emotional memory processes.^20,21^ Recent evidence suggests that AMY-HPC interactions are pivotal to non-emotional WM tasks, and memory load can be decoded by the information flow from the HPC to the AMY in the low-frequency band during maintenance.^22^ Notably, the HPC is not a unitary structure, and the interconnections between the AMY and HPC subregions are complex.^23^ The HPC can be divided into anterior and posterior parts (aHPC and pHPC), corresponding to the ventral and dorsal HPC in rodents, which exhibit differences in function, structure, and connections to cortical and subcortical areas.^24^ It remains unclear whether the connectivity between the AMY and distinct HPC subregions exhibits different patterns during WM maintenance. Intracranial electroencephalography (iEEG) is a novel technique that allows for simultaneous measurements from multiple recording sites of interest.

To investigate the role of the AMY and AMY-HPC interactions during WM maintenance, we analyzed iEEG recordings from the AMY (including both the aHPC and pHPC) and HPC in nine epileptic patients.^25^ A modified Sternberg paradigm was implemented, and the patients performed a verbal WM task that included trials of different set sizes. WM load was determined by the number of letters. Our results showed increased low-frequency power and intraregional theta‒gamma phase-amplitude coupling (PAC) under the high-load condition in both the AMY and HPC during maintenance. At the network level, a high WM load elicited an increase in the modulation strength of HPC (including both the aHPC and pHPC) theta phase entraining AMY gamma amplitude. Additionally, the aHPC and pHPC exhibited divergent connectivity patterns with the AMY, was characterized by increased AMY-aHPC theta phase synchrony and directional connectivity strength, with the influence direction being from the aHPC to the AMY, under the high-load condition, while pHPC-AMY theta phase synchrony was not affected by memory load. These load-dependent connectivity characteristics significantly contribute to behavioral success.

## 2. Results

### 2.1. Task, behavior and electrode locations

Nine epilepsy patients (5 females and 4 males) participated in a modified Sternberg WM task (Fig. 1a). All the subjects performed well on the task, with a correct response rate of 92.3 ± 2.7% (for both the IN and OUT trials across sets with sizes of 4, 6 and 8). As illustrated in the original work, the mean memory capacity was 6.8 in this dataset,^25^ reflecting the limited number of objects that could be held in the WM. Given that the interregional coupling pattern varied between conditions below and above the capacity limit,^26^ in our study, low and high WM loads refer to trials with set sizes of 4 and 6, both of which were under the capacity limit. Focusing on trials within the capacity limit avoids introducing potential confounding factors that exceed the memory capacity. The subjects achieved greater accuracy under the low-load condition (98.7 ± 1.6% correct responses) than under the high-load condition (91.3 ± 4.4% correct responses) (t (8) = 5.717, p = 0.0004, paired t test). Across all subjects, the response time (RT) for correct trials significantly increased with WM load from a set size of 4 to 6 (1.2 ± 0.2 s versus 1.4 ± 0.2 s, respectively) (t (8) = 4.664, p = 0.0016, paired t test) (Fig. 1b). Both behavioral measures (accuracy, RT) were load dependent, exhibiting better performance under the low-load condition.

**Fig. 1.**
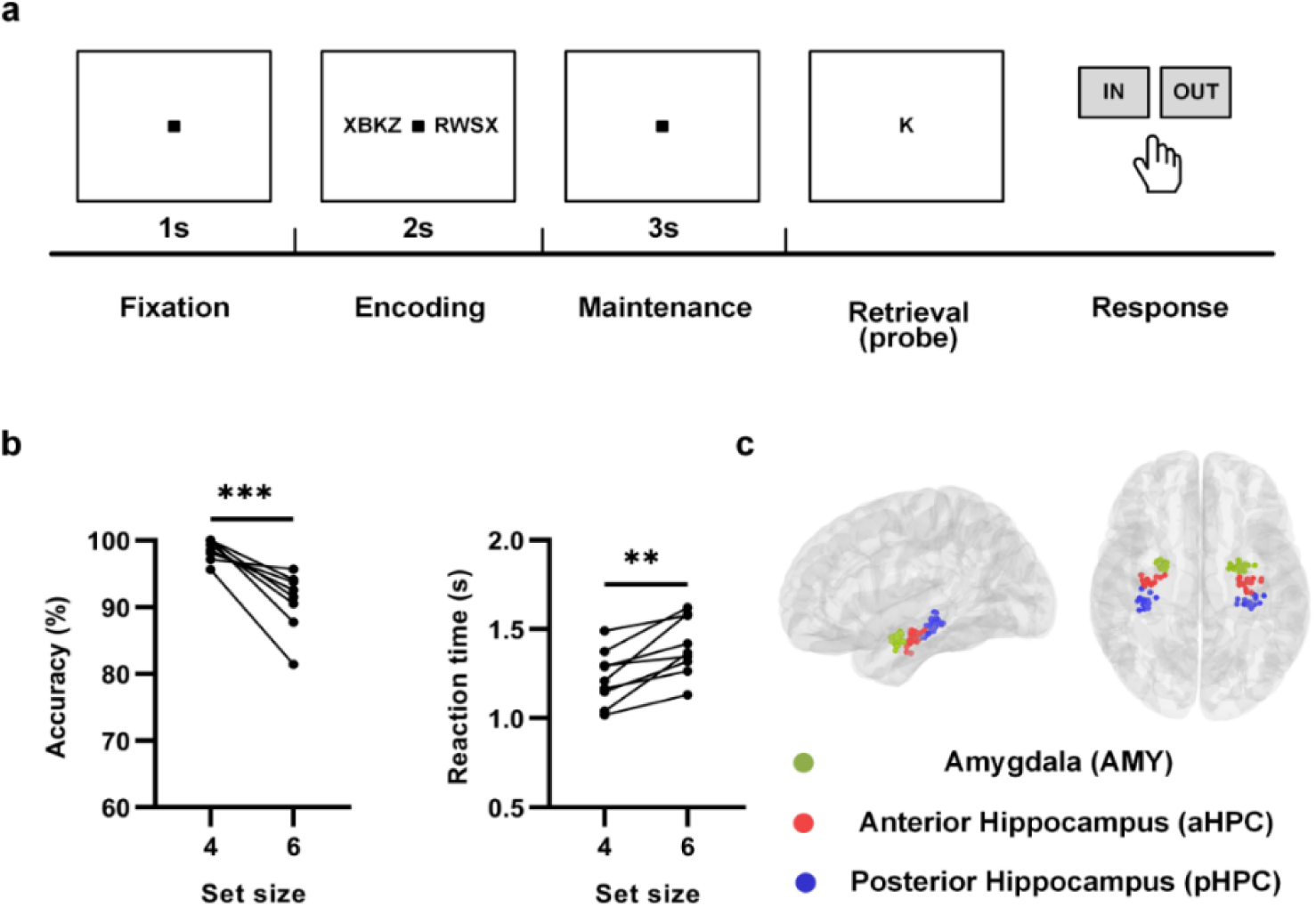
Task, behavior results and electrode locations. **(a)** Experimental paradigm. Subjects performed a modified Sternberg task, which incorporated temporally separated epochs of the encoding, maintenance and retrieval phases of WM. Each trial started with a blank screen with a fixation square for 1 second (fixation period), followed by the presentation of a set of 4, 6, or 8 letters in the center of the screen surrounded by ‘X’. The letter ‘X’ was not part of the set to be memorized. The stimulus was presented for 2 seconds (encoding period). Following the encoding period, the stimulus disappeared from the screen and was replaced by a fixation square for 3 seconds (maintenance period). After the maintenance period, a probe letter appeared. Subjects were instructed to respond rapidly to indicate whether the probe letter was part of the stimulus set by pressing a button (“IN” or “OUT”). Each session consisted of 50 trials lasting approximately 10 minutes. Each subject completed 2−7 sessions. **(b)** Accuracy decreased (t (8) = 5.717, p = 0.0004, paired t test) and RT increased (t (8) = 4.664, p = 0.0016, paired t test) from set size 4 to set size 6. Dots indicate individual results. ***p<0.001, **p<0.01. **(c)** Electrode locations across 9 subjects in the MNI space. Recording regions are marked using different colors (green, AMY; red, aHPC; blue, pHPC).

Across 9 patients, the two most medial channels on each electrode targeting each region in each hemisphere were selected. We analyzed iEEG data from a total of 30, 32, and 32 electrodes in the AMY, aHPC, and pHPC, respectively (Fig. 1c). All analyses in this study were conducted after bipolar-referenced processing.

### 2.2. Load-dependent low-frequency power in the AMY during WM maintenance

To investigate the influence of WM load on AMY activity during maintenance, we analyzed the low-frequency (3–8 Hz theta range and 8–13 Hz alpha range) power responses of the AMY in the correct trials under both low- and high-load conditions. The time-frequency results indicated significant load effects on clusters in the AMY, as well as in the aHPC and pHPC, with a notable increase in low-frequency power for the high-load condition (cluster-based permutation, p < 0.05, Fig. 2a, b, c).

**Fig. 2.**
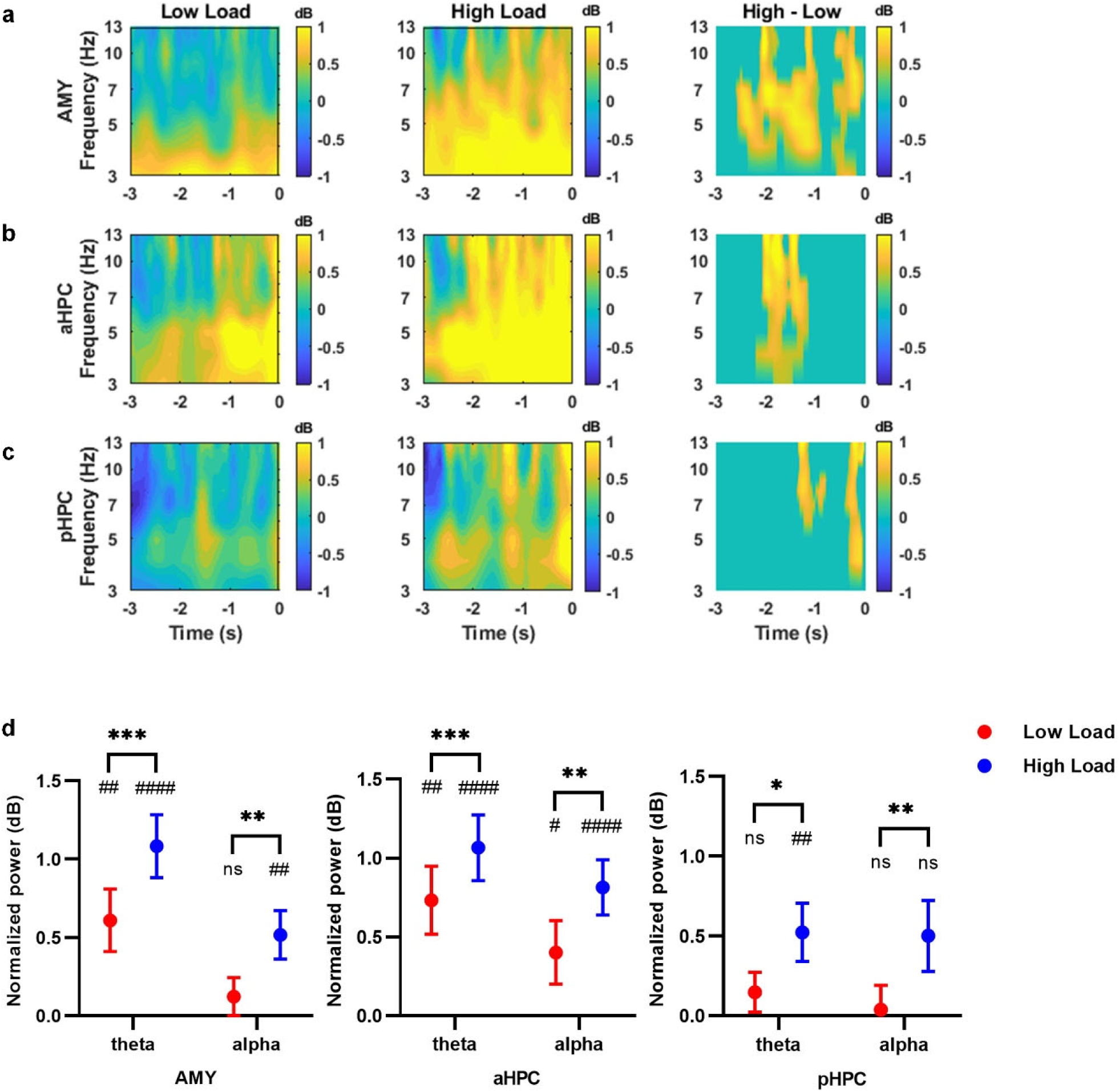
Low-frequency power during maintenance for the correct trials. Normalized power in the **(a)** AMY (electrodes N = 30), **(b)** aHPC (electrodes N = 32) and **(c)** pHPC (electrodes N = 32) areas. Left and middle: time-frequency spectrograms for low- and high-load trials. Right: difference map (high - low) with significant clusters after cluster-based permutation testing (n = 2000 permutations). Time ‘0’ indicates the onset of the retrieval period. **(d)** Averaged theta (3–8 Hz) and alpha (8–13 Hz) power for low- and high-load trials. Wilcoxon signed-rank test (two-tailed) showed that theta and alpha power were increased significantly for high-load trials compared to low-load trials in the AMY (theta: p = 0.0003, alpha: p = 0.0028), aHPC (theta: p = 0.0006, alpha: p = 0.0012) and pHPC (theta: p = 0.0358, alpha: p = 0.0013). One sample Wilcoxon test showed that significant increases in theta power during maintenance relative to baseline under both low- and high-load conditions in the AMY (low load: p = 0.0011, high load: p = 1.6838e-06) and aHPC (low load: p = 0.0093, high load: p = 1.2635e-05), while significant power increases in the pHPC were observed only under the high-load condition (p = 0.0088). One sample Wilcoxon test also showed that significant increases in alpha power during maintenance relative to baseline under the high-load condition in the AMY (p = 0.0035), and under both low- and high-load conditions in the aHPC (low load: p = 0.0454, high load: p = 6.6853e-05;). Error bars represent standard errors of the mean (SEMs) across all electrodes. Symbols represent statistically significant differences: * compared to different loads; # compared to baseline. ***p<0.001, **p<0.01, *p<0.05; ####p<0.0001, ##p<0.01, #p<0.05.

We subsequently extracted and averaged the power within the theta and alpha bands and then compared these powers between the low- and high-load conditions. High loads resulted in increased theta and alpha power (Fig. 2d) in all three brain regions (aHPC, theta: p = 0.0006; alpha: p = 0.0012; pHPC, theta: p = 0.0358; alpha: p = 0.0013; AMY, theta: p = 0.0003; alpha: p = 0.0028; Wilcoxon signed-rank test, two-tailed; AMY electrodes N = 30, aHPC electrodes N = 32, pHPC electrodes N = 32). The increased low-frequency power of the HPC under the high-load condition was consistent with previous studies.^10,11^

To validate that the observed power changes were induced by the WM task, we compared the power during maintenance with that at baseline and found an increase in low-frequency power in the AMY, aHPC and pHPC (Fig. 2d), especially under the high-load condition. Specifically, the AMY and aHPC exhibited task-induced power increases in the theta band under both load conditions (AMY low load: p = 0.0011, high load: p = 1.6838e-06; aHPC low load: p = 0.0093, high load: p = 1.2635e-05; one sample Wilcoxon test), while significant power changes in the pHPC were observed only under the high-load condition (low load: p = 0.1966; high load: p = 0.0088; one sample Wilcoxon test). In the alpha band, only the aHPC exhibited a significant increase in power under both load conditions (low load: p = 0.0454; high load: p = 6.6853e-05; one sample Wilcoxon test). A significant increase in alpha power in the AMY was observed exclusively under the high-load condition (low load: p = 0.1772; high load: p = 0.0035; one sample Wilcoxon test), whereas no task-induced power changes in the pHPC were detected under either load condition (low load: p = 0.7053; high load: p = 0.0521; one sample Wilcoxon test). Taken together, these results suggest that not only is the HPC (including aHPC and pHPC) involved in maintenance and exhibits load dependency but the AMY is also recruited, with its activity being affected by WM load in a manner similar to that of the HPC. Notably, these three regions exhibited load dependency within different temporal ranges on the time-frequency spectrum, suggesting that they may have distinct roles in WM maintenance.

### 2.3. Greater theta-gamma phase-amplitude coupling in the AMY under the high-load condition during WM maintenance

It has been suggested that the organization of WM for multiple items is facilitated by theta‒gamma coupling in the HPC during maintenance.^9,27^ We wondered whether the neural mechanism of theta‒ gamma coupling in the AMY could coordinate WM maintenance. We calculated theta (3–8 Hz)-gamma (30–100 Hz) PAC values for the correct trials during maintenance under different load conditions. Then, the raw PAC was z-scored against a null distribution. The theta phase entrained the gamma amplitude in the AMY (low-load: peaks at 7 Hz and 73 Hz for phase and amplitude frequencies, respectively; high-load: peaks at 8 Hz and 78 Hz for phase and amplitude frequencies, respectively; Fig. 3a left), aHPC (low-load: peaks at 7 Hz and 98 Hz for phase and amplitude frequencies, respectively; high-load: peaks at 5 Hz and 93 Hz for phase and amplitude frequencies, respectively; Fig. 3a middle) and pHPC (low-load: peaks at 5 Hz and 98 Hz for phase and amplitude frequencies, respectively; high-load: peaks at 4 Hz and 98 Hz for phase and amplitude frequencies, respectively; Fig. 3a right).

**Fig. 3.**
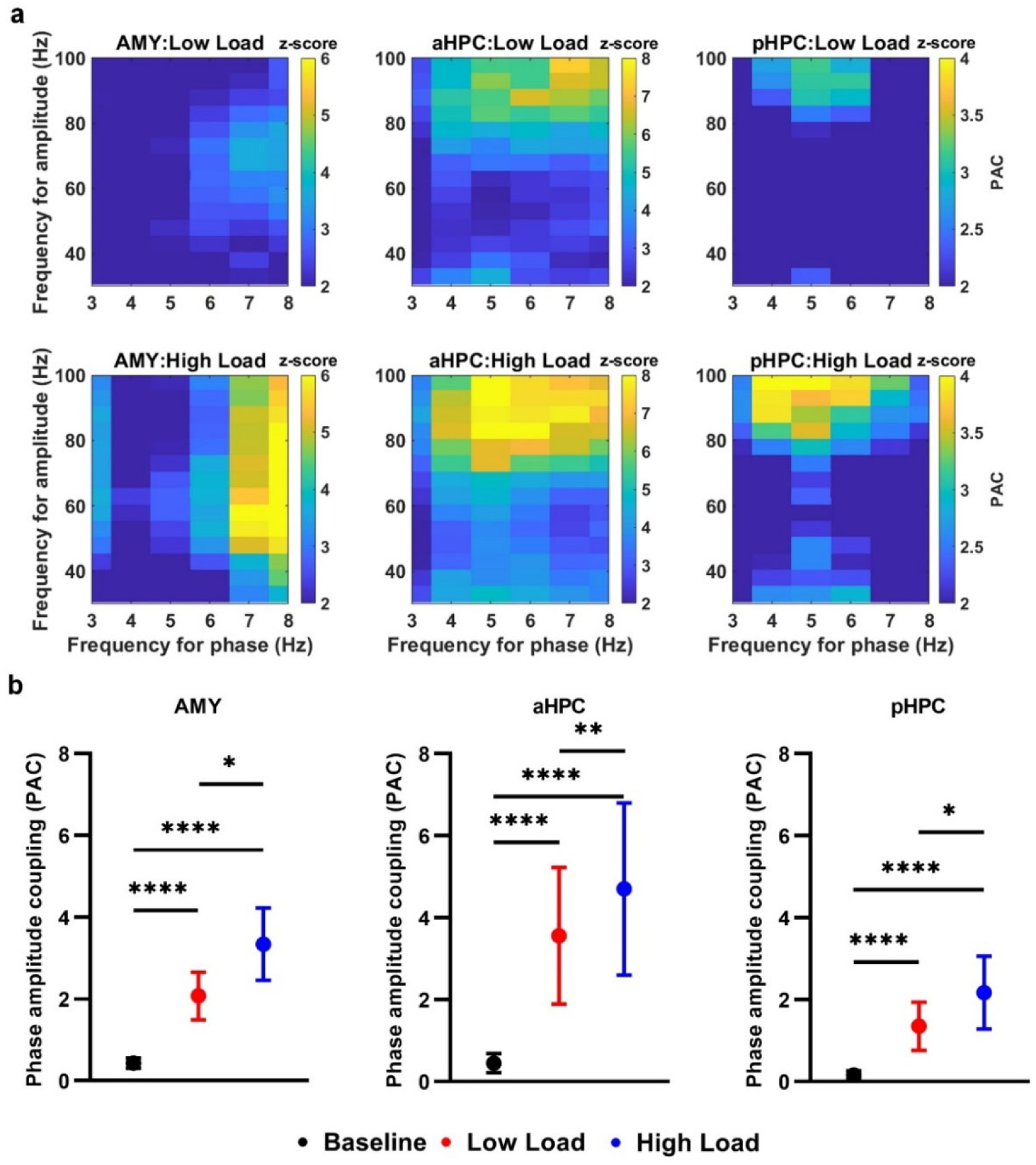
Intraregional theta-gamma PAC for the correct trials. **(a)** Averaged phase-amplitude comodulogram during maintenance in the AMY (left, electrodes N = 29), aHPC (middle, electrodes N = 32) and pHPC (right, electrodes N = 30) under both low- and high-load conditions. Warmer colors indicate stronger modulation. **(b)** Wilcoxon signed-rank test (two-tailed) showed that average theta (3–8 Hz)-gamma (30–100 Hz) PAC was significantly enhanced during maintenance compared to baseline in the AMY, aHPC and pHPC under both load conditions (AMY: low-load p = 1.1928e-05, high-load p = 3.6284e-05; aHPC: low-load p = 1.1437e-06, high-load p = 1.6019e-07; pHPC: low-load p = 8.8565e-05, high-load p = 9.9838e-07). Wilcoxon signed-rank test (two-tailed) also showed that theta-gamma PAC was significantly greater under the high-load condition than under the low-load condition in the AMY (p = 0.048), aHPC (p = 0.0015) and pHPC (p = 0.0473). Error bars represent the SEMs across electrodes. ****p<0.0001, **p<0.01, *p<0.05.

We assessed task-induced changes in the strength of the intraregional PAC during maintenance compared to baseline. Across all loads, we found that the pattern of the intraregional PACs in the three brain regions during maintenance increased relative to that at baseline (AMY: low-load p = 1.1928e-05, high-load p = 3.6284e-05; aHPC: low-load p = 1.1437e-06, high-load p = 1.6019e-07; pHPC: low-load p = 8.8565e-05, high-load p = 9.9838e-07; Wilcoxon signed-rank test, two-tailed; AMY electrodes N = 29, aHPC electrodes N = 32, pHPC electrodes N = 30; Fig. 3b). Then, we compare the intraregional PAC during maintenance between different load conditions. The AMY exhibited a significant enhancement in theta-gamma PAC under the high-load condition compared to the low-load condition (p = 0.048, Wilcoxon signed-rank test, two-tailed). Similar to a previous study,^9^ our results also showed that simultaneous maintenance of multiple items in WM is accompanied by theta‒gamma coupling in the HPC. We found significantly greater intraregional theta-gamma PAC value s under the high-load condition for both the aHPC and pHPC (aHPC: p = 0.0015; pHPC: p = 0.0473; Wilcoxon signed-rank test, two-tailed; Fig. 3b). Taken together, our results confirmed the engagement of the AMY in the maintenance process and demonstrated that WM load enhanced the interregional theta-gamma PAC for the AMY.

### 2.4. Load-dependent AMY-aHPC theta synchrony during WM maintenance

Considering the supposed role of low-frequency oscillations in interregional communication^28^, we investigated the phase synchrony between the AMY and HPC for the correct trials during maintenance under different load conditions. The weighted phase lag index (wPLI) serves as a conservative measure of phase synchrony, offering the advantages of low sensitivity to volume conduction and uncorrelated noise sources.^29^

As shown in Fig. 4a, b, the time-frequency domain wPLI revealed dynamic fluctuations in phase synchrony between each pair of regions in specific frequency ranges during maintenance. We first tested whether maintenance periods exhibited task-induced phase synchrony. We quantitatively averaged the theta band (from 3 to 8 Hz) wPLI and found that phase synchrony during maintenance exhibited significant task-induced enhancement under the high-load condition for both the AMY-aHPC and AMY-pHPC (p = 0.0007 and p = 1.7927e-05 for the AMY-aHPC and AMY-pHPC, respectively; Wilcoxon signed-rank test, two-tailed; AMY-aHPC electrode pairs N = 48, AMY-pHPC electrode pairs N = 33, Fig. 4b). However, AMY-aHPC theta phase synchrony was not significantly greater for the maintenance phase than at baseline under the low-load condition, while the AMY-pHPC showed task-induced theta phase synchrony (p = 0.2427 and p = 0.0022 for the AMY-aHPC and AMY-pHPC, respectively; Wilcoxon signed-rank test, two-tailed, Fig. 4b). Next, we explored the effect of WM load on theta phase synchrony during maintenance. The results showed a significant load effect in the AMY-aHPC with increasing wPLI for the high-load condition relative to the low-load condition (p = 0.0053, Wilcoxon signed-rank test, two-tailed, Fig. 4b). However, no significant difference in the theta band wPLI of the AMY-pHPC under different load conditions was detected (p = 0.1827, Wilcoxon signed-rank test, two-tailed, Fig. 4b). Additionally, we evaluated the phase synchrony in the alpha band (from 8 to 13 Hz). No significant difference between baseline and maintenance was observed for either the AMY-aHPC (baseline vs. low-load maintenance: p = 0.9070, baseline vs. high-load maintenance: p = 0.1963; Wilcoxon signed-rank test, two-tailed) or AMY-pHPC (baseline vs. low-load maintenance: p = 0.3483, baseline vs. high-load maintenance: p = 0.5135; Wilcoxon signed-rank test, two-tailed), regardless of the load condition (Fig. 4c). Although we observed that the alpha phase synchrony between the AMY and pHPC was not significantly greater during maintenance than at baseline, a high load resulted in greater AMY-pHPC alpha phase synchrony (p = 0.0443, Wilcoxon signed-rank test, two-tailed, Fig. 4c). In contrast, phase synchrony between the AMY and aHPC showed no difference in the alpha band between different load conditions (p = 0.084). These results provide clear evidence that multi-item WM maintenance is associated with increased theta phase synchrony between the AMY and HPC. Moreover, we found that WM load had divergent effects on the connectivity patterns between the AMY and HPC subregions.

**Fig. 4.**
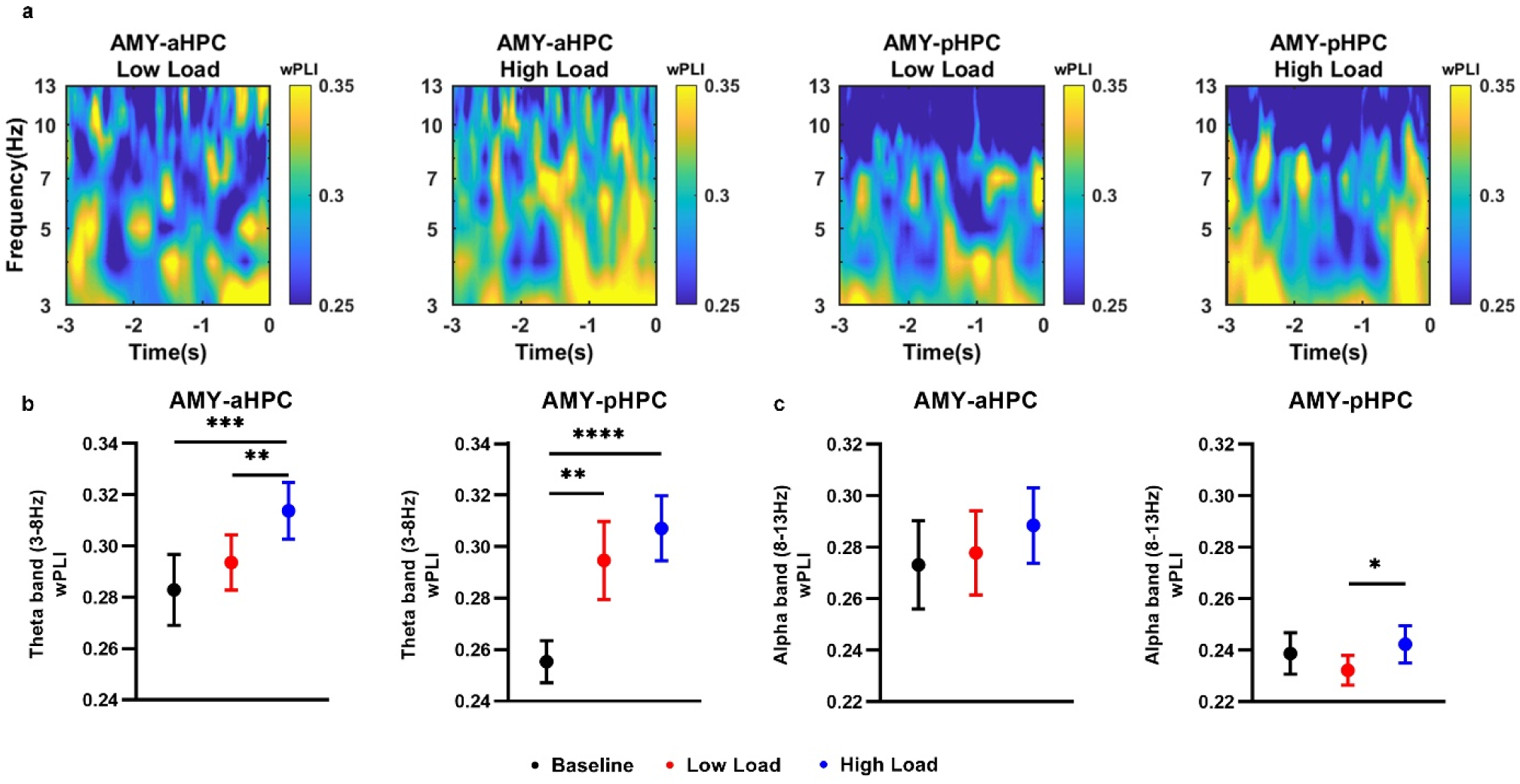
Interregional phase synchrony between the AMY and aHPC/pHPC for the correct trials in the low-frequency band. **(a)** Phase synchrony (i.e., wPLI) over the AMY-aHPC and AMY-pHPC under both low- and high-load conditions (AMY-aHPC electrode pairs N = 48, AMY-pHPC electrode pairs N = 33). The wPLI value ranges from 0 to 1, with warmer colors indicating greater phase synchrony. Time ‘0’ indicates the onset of the retrieval period. **(b, c)** Averaged theta (3–8 Hz) and alpha (8–13 Hz) wPLI values during baseline and during low- and high-load maintenance. Under the high-load condition, theta band wPLI values over the AMY-aHPC and AMY-pHPC were significantly enhanced during maintenance compared to baseline (p = 0.0007 and p = 1.7927e-05, respectively). Under the low-load condition, theta band wPLI values over the AMY-pHPC were significantly greater during maintenance compared to baseline (p = 0.0022). Over the AMY-aHPC, theta band wPLI was significantly greater under the high-load condition compared to the low-load condition (p = 0.0053). Over the AMY-pHPC, alpha band wPLI was significantly greater under the high-load condition compared to the low-load condition (p = 0.0443). Error bars indicate the SEMs across electrode pairs. ****p<0.0001, ***p<0.001, **p<0.01, *p<0.01, indicate significant differences using paired Wilcoxon signed-rank test (two-tailed).

### 2.5. WM load effects on the directional interaction between the AMY and HPC in the theta band

In addition to using wPLI measurements, which reveal only non-directional interactions, we utilized frequency-domain Granger causality (GC) analysis to further elucidate the directionality of information transfer in the theta band between different brain regions. We found that the GC index under the high-load condition was significantly greater than that under the low-load condition only in the aHPC-to-AMY direction (AMY-to-aHPC p = 0.9001, aHPC-to-AMY p = 0.008, AMY-to-pHPC p = 0.1736, pHPC-to-AMY p = 0.8793; Wilcoxon signed-rank test, two-tailed; AMY-aHPC electrode pairs N = 43, AMY-pHPC electrode pairs N = 34; Fig. 5a). Then, we estimated the directionality of the net information flow over the aHPC-AMY and pHPC-AMY circuits. The results of the net information flow analysis, ΔGranger (GC index of aHPC-to-AMY minus GC index of AMY-to-aHPC), revealed that the directional influence in the theta band from aHPC to AMY was significantly greater under the high-load condition (p = 0.0423, Wilcoxon signed-rank test, two-tailed; Fig. 5b), while there was no significant directional preference between the AMY and pHPC (aHPC-AMY p = 0.0423, pHPC-AMY p = 0.5542). Taken together, our findings indicated that the theta-driven directional influence from the aHPC to the AMY was highly load dependent, exhibiting increased causal connectivity under the high-load condition. This observation is consistent with the phase synchrony results, indicating that the WM load regulates the connectivity strength between the AMY and aHPC in the theta band.

**Fig. 5.**
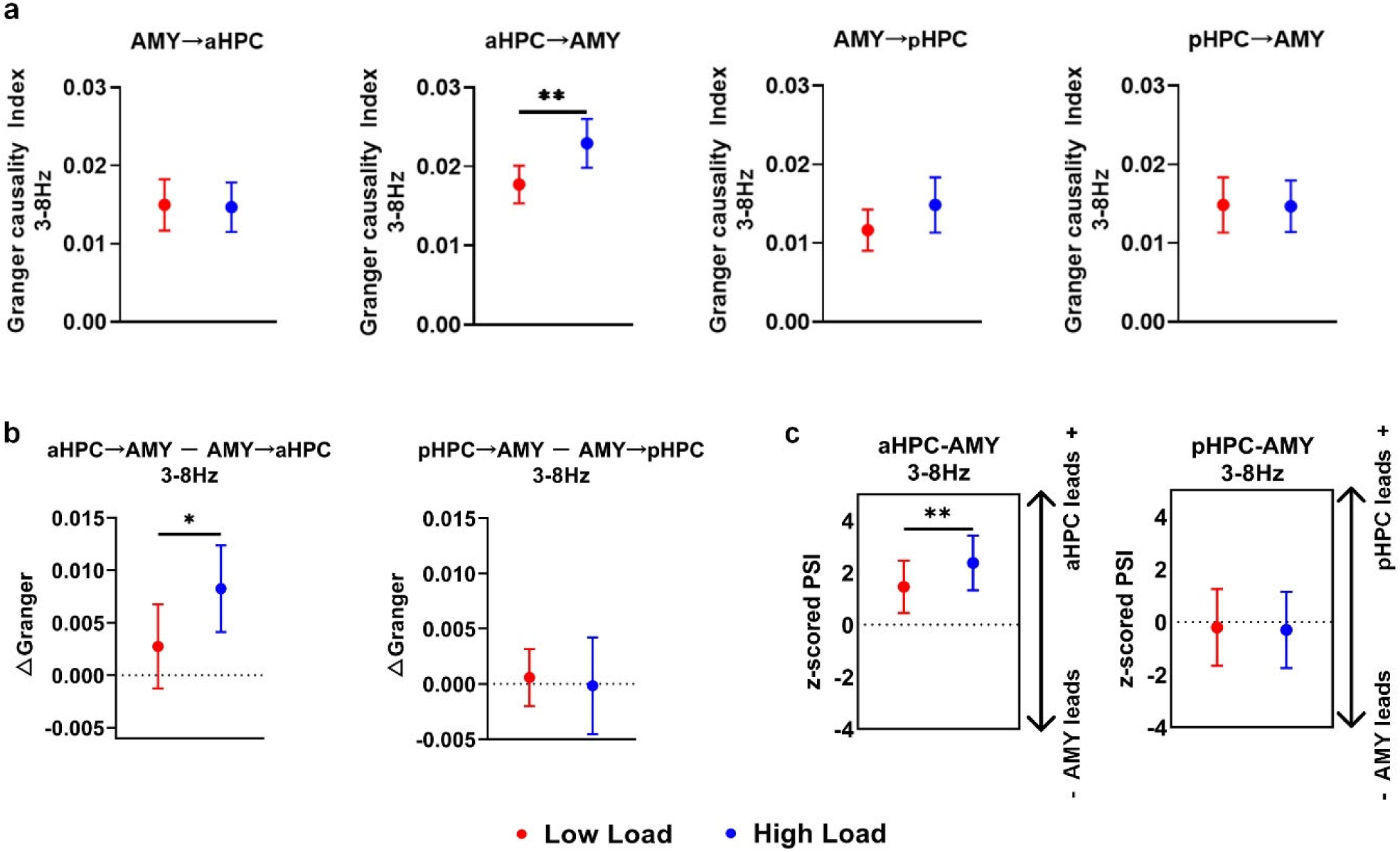
Directional information flow in the theta band between the AMY and aHPC/pHPC during maintenance for the correct trials. **(a)** GC index in the aHPC-to-AMY direction was increased significantly under the high-load condition compared to the low-load condition (p = 0.008; N = 43). **(b)** Net information flow (ΔGranger) of aHPC-to-pHPC (GC index of aHPC-to-AMY minus GC index of AMY-to-aHPC) was increased under the high-load condition compared to the low-load condition (p = 0.0423). **(c)** The z-scored PSI value. Positive values indicate that the aHPC (left) or pHPC (right) transfers to the AMY, and negative values indicate that the AMY transfers to the aHPC or pHPC (significance was thresholded at |z| > 1.96). PSI of the aHPC leads AMY connectivity was greater under the high-load condition compared to the low-load condition (high vs. low: mean z-scored PSI 2.3687 vs. 1.4547, p = 0.0092). Error bars represent the SEMs across electrode pairs. **p<0.01, *p<0.05, indicate significant differences using paired Wilcoxon signed-rank test (two-tailed).

Additionally, we re-assessed the information flow direction in the theta band between the AMY and aHPC/pHPC by employing the phase slope index (PSI), which is more insensitive to mixtures of independent sources than GC.^30^ We found that the load-dependent directional influence was specific to the aHPC-AMY circuit, characterized by significantly stronger unidirectional aHPC-driven connectivity under the high-load condition (high vs. low: mean z-scored PSI 2.3687 vs. 1.4547, p = 0.0092, Wilcoxon signed-rank test, two-tailed; AMY-aHPC electrode pairs N = 16, AMY-pHPC electrode pairs N = 8; Fig. 5c left). However, no significant directional influence within the pHPC-AMY circuit was observed in either the low- or high-load trials (high vs. low: mean z-scored PSI - 0.3061 vs. −0.209, p = 0.7422; Fig. 5c right). This result was consistent with the results from the above wPLI and GC analyses showing that aHPC-AMY theta phase synchrony was load dependent, with a high load eliciting stronger directional connectivity from the aHPC to the AMY.

### 2.6. Load-dependent interregional phase-amplitude coupling between the AMY and HPC during WM maintenance

Given that coupled theta and gamma activity over the AMY-HPC circuit tracked successful subsequent remembered responses to emotional stimuli,^31^ we examined whether interregional PAC between the AMY and HPC was associated with successful non-emotional WM maintenance. PAC plays a crucial role in the flexible coordination of interregional information transfer, whereby the phase of slow oscillations dynamically adjusts the magnitude of high-frequency amplitude.^32^ We performed interregional PAC analyses of both phase-amplitude combinations (low-frequency phase from AMY and high-frequency amplitude from aHPC or pHPC, and vice versa) for the correct trials. The z-scored PAC showed that modulations of the theta phase of the AMY entrained the gamma amplitude of the aHPC/pHPC (AMY-aHPC: peak at 4 Hz and 38 Hz under the low-load condition, 3 Hz and 38 Hz under the high-load condition; AMY-pHPC: peak at 4 Hz and 43 Hz under the low-load condition, 3 Hz and 98 Hz under the high-load condition); and the theta phase of the aHPC/pHPC entrained the gamma amplitude of the AMY (aHPC-AMY: peak at 6 Hz and 78 Hz under the low-load condition, 6 Hz and 88 Hz under the high-load condition; pHPC-AMY: peak at 7 Hz and 63 Hz under the low-load condition, 7 Hz and 53 Hz under the high-load condition) (Fig. 6a).

**Fig. 6.**
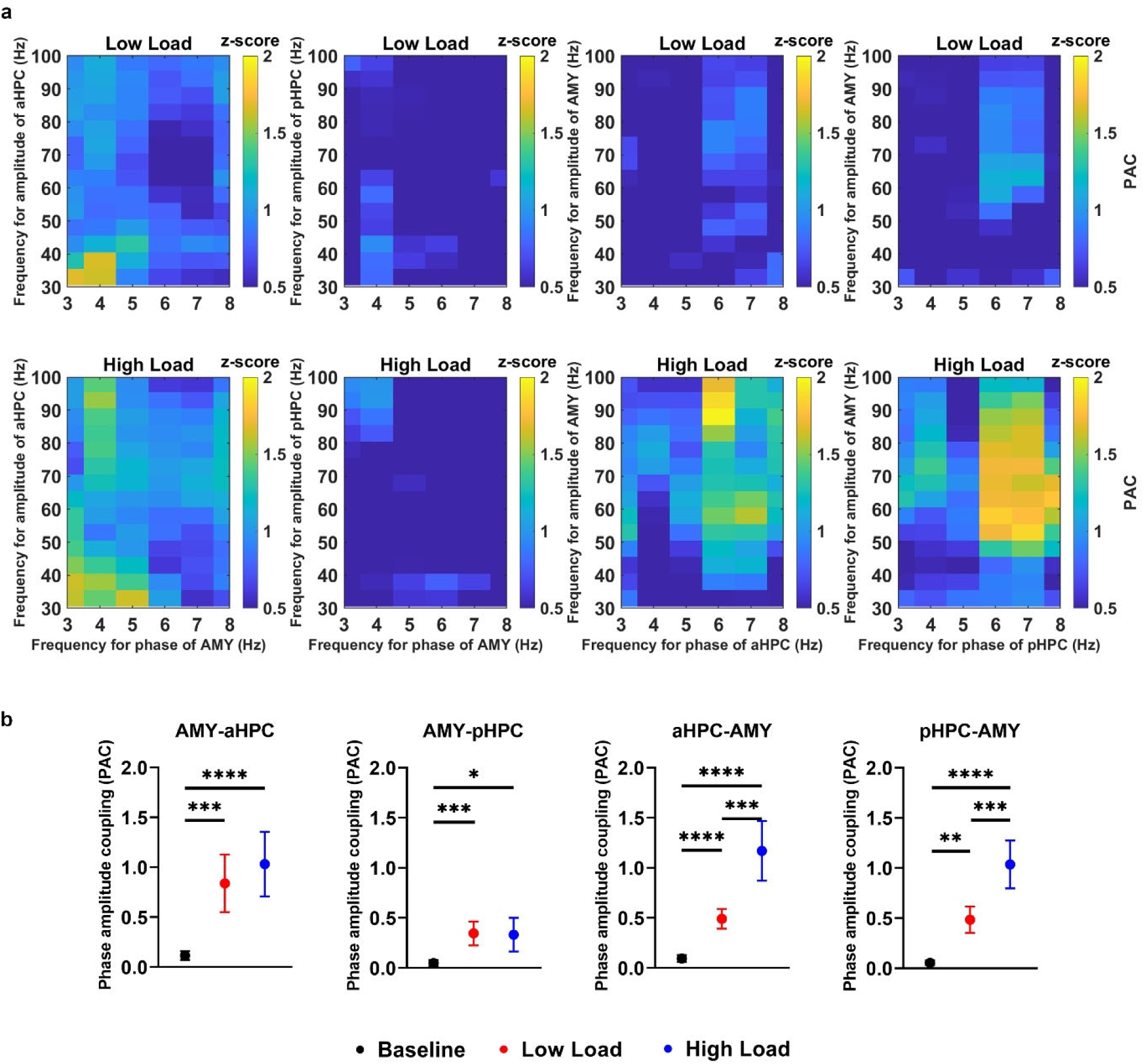
Interregional theta-gamma PAC between the AMY and aHPC/pHPC for the correct trials. **(a)** Averaged phase-amplitude comodulogram across electrode pairs during maintenance. Warmer colors denote higher z-scored PAC values. **(b)** Averaged interregional theta (3–8 Hz)-gamma (30–100 Hz) PAC during baseline, low-load and high-load maintenance. ‘AMY-aHPC’ denotes the AMY theta phase entraining aHPC gamma amplitude, while ‘aHPC-AMY’ indicates the reverse modulation. Theta-gamma PAC values of the AMY-aHPC, AMY-pHPC, aHPC-AMY and pHPC-AMY were significantly increased during maintenance compared to baseline under both low- (AMY-aHPC: p = 1.0567e-04; AMY-pHPC: p = 7.5029e-04; aHPC-AMY: p = 1.1955e-06; pHPC-AMY: p = 0.0067) and high-load (AMY-aHPC: p = 6.1486e-07; AMY-pHPC: p = 0.0137; aHPC-AMY: p = 3.8502e-07; pHPC-AMY: p = 4.0286e-06) conditions. aHPC and pHPC phase of theta frequency to AMY gamma amplitude coupling are significantly greater under the high-load condition compared to the low-load condition (aHPC-AMY: p = 8.9365e-04, pHPC-AMY: p = 2.2114e-04). Error bars represent the SEMs across electrode pairs. ****p<0.0001, ***p<0.001, **p<0.01, indicate significant differences using paired Wilcoxon signed-rank test (two-tailed).

We then evaluated whether the interregional PAC was task induced. Both AMY-aHPC/pHPC and aHPC/pHPC-AMY exhibited task-specific cross-frequency coupling under both low-load (AMY-aHPC: p = 1.0567e-04; AMY-pHPC: p = 7.5029e-04; aHPC-AMY: p = 1.1955e-06; pHPC-AMY: p

= 0.0067) and high-load conditions (AMY-aHPC: p = 6.1486e-07; AMY-pHPC: p = 0.0137; aHPC-AMY: p = 3.8502e-07; pHPC-AMY: p = 4.0286e-06, Wilcoxon signed-rank test, two-tailed; AMY-aHPC electrode pairs N = 56, aHPC-AMY electrode pairs N = 54, AMY-pHPC electrode pairs N = 51, pHPC-AMY electrode pairs N = 50) (Fig. 6b). To further compare the PAC between the two load conditions, we calculated the average theta‒gamma PAC and observed a significantly greater PAC under the high-load condition for aHPC–AMY and pHPC–AMY (p = 8.9365e-04, p = 2.2114e-04, Wilcoxon signed-rank test, two-tailed) (Fig. 6b). These results suggest that interregional PAC between the HPC and AMY supports the process of WM in a load-dependent manner, with the phase of HPC theta oscillations modulating the gamma amplitude of the AMY during maintenance.

### 2.7. Absence of AMY-HPC interactions in the incorrect trials

Considering that the maintenance process involves keeping novel information available and contributes significantly to successful behavioral performance, we investigated the specificity of the above findings to successful outcomes. We first analyzed the power change between baseline and maintenance for the incorrect trials. In the theta/alpha band, only the AMY exhibited a significant task-induced increase in power during maintenance (p = 0.012 and p = 0.011, one sample Wilcoxon test), whereas the aHPC and pHPC showed no such change (theta band: p = 0.9046 and p = 0.7609; alpha band: p = 0.1838 and p = 0.8034, one sample Wilcoxon test) (Fig. 7a). Next, we assessed the theta band wPLI for the AMY-aHPC and AMY-pHPC circuits for the incorrect trials; this characteristic indicated task relevance and load dependency in the preceding analyses. No significant difference between maintenance and baseline was found (p = 0.6291 and p = 0.9632 for the AMY-aHPC and AMY-pHPC, respectively, Wilcoxon signed-rank test, two-tailed; Fig. 7b).

**Fig. 7.**
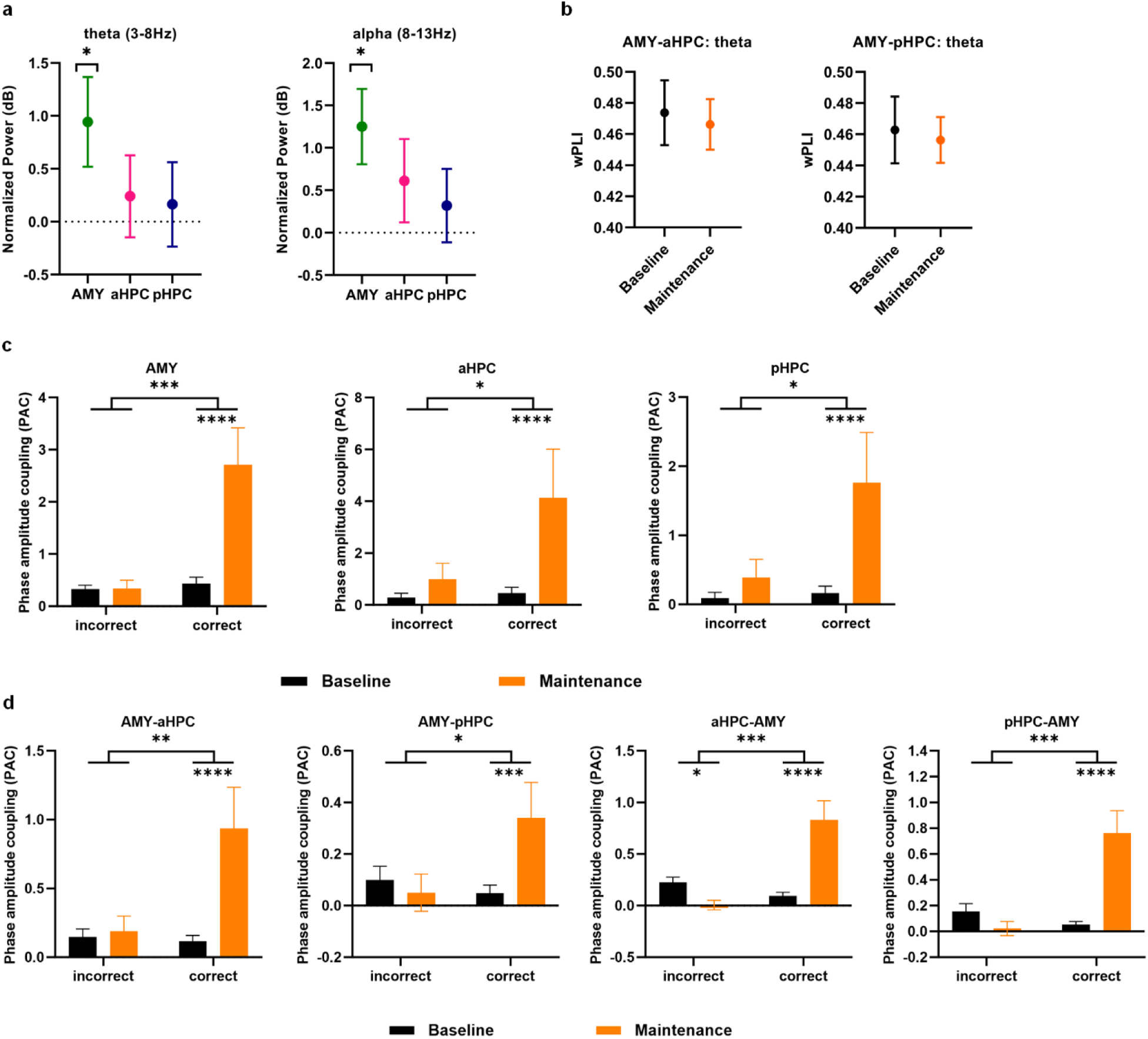
Intraregional local activity and AMY-aHPC/pHPC interactions for the incorrect trials. **(a)** One sample Wilcoxon test showed that theta (3–8 Hz) and alpha (8–13 Hz) power were increased in the AMY during maintenance compared to baseline (theta power: p = 0.012, alpha power: p = 0.011). **(b)** AMY-aHPC/pHPC wPLI within the theta band. Wilcoxon signed-rank test (two-tailed) showed that there was no significant difference between baseline and maintenance over AMY-aHPC (p = 0.6291) and AMY-pHPC (p = 0.9632). **(c)** Intraregional theta (3–8 Hz)-gamma (30–100 Hz) PAC. Repeated-measures ANOVA showed the intraregional PAC changes from baseline to maintenance were associated with task performance (correct/incorrect) in the AMY (p = 0.0008, F(1,28) = 13.945), aHPC (p = 0.0252, F(1,31) = 5.5309) and pHPC (p = 0.01, F(1,29) = 7.5957). Wilcoxon signed-rank test (two-tailed) indicated that intraregional PAC was significantly enhanced during maintenance compared to baseline for the correct trials (AMY: p = 5.0274e-05; aHPC: p = 4.6223e-06; pHPC: p = 3.1123e-05), but not for the incorrect trials (AMY: p = 0.5096, aHPC: p = 0.3309, pHPC: p = 0.2059). **(d)** Interregional theta-gamma PAC over AMY-aHPC/pHPC (AMY theta phase entraining aHPC/pHPC gamma amplitude) and aHPC/pHPC-AMY (aHPC/pHPC theta phase entraining AMY gamma amplitude). The interregional PAC in both AMY-aHPC/pHPC and aHPC/pHPC-AMY showed significant interaction effects of task performance (correct/incorrect) and stage (baseline/maintenance) (AMY-aHPC: p = 0.0018, F (1, 55) = 10.6819; AMY-pHPC: p = 0.0129, F (1, 50) = 6.6488; aHPC-AMY: p = 0.0001, F (1, 53) = 16.7314; pHPC-AMY: p = 0.0001, F (1, 49) = 17.2807; repeated-measures ANOVA). For the incorrect trials, Wilcoxon signed-rank test (two-tailed) revealed no significant difference between baseline and maintenance for the AMY-aHPC (p = 0.7197), AMY-pHPC (p = 0.8882), and pHPC-AMY (p = 0.6055); the aHPC-AMY was lower during maintenance compared to baseline (p = 0.0161). However, for the correct trials, the PAC values were significantly greater during maintenance for both AMY-aHPC/pHPC (AMY-aHPC: p = 2.621e-06, AMY-pHPC: p = 5.4257e-04) and aHPC/pHPC-AMY (aHPC-AMY: p = 6.6129e-07, pHPC-AMY: p = 4.1707e-05). Error bars represent the SEMs across electrodes (power and intraregional PAC) or electrode pairs (wPLI and interregional PAC). ****p<0.0001, ***p<0.001, **p<0.01, *p < 0.05.

We then quantified the intraregional theta‒gamma PAC at baseline and maintenance. A 2 (performance: incorrect vs. correct) × 2 (stage: baseline vs. maintenance) repeated-measures ANOVA revealed significant interaction effects in all three regions (AMY: p = 0.0008, F (1,28) = 13.945; aHPC: p = 0.0252, F (1,31) = 5.5309; pHPC: p = 0.01, F (1,29) = 7.5957), indicating that task performance was associated with the PAC changes from baseline to maintenance in AMY, aHPC and pHPC. Post-hoc pairwise comparisons using two-tailed Wilcoxon signed-rank test showed distinct patterns between performance conditions. For the incorrect trials, PAC values during maintenance did not significantly differ from baseline in the AMY (p = 0.5096), aHPC (p = 0.3309) and pHPC (p = 0.2059). In contrast, correct trials demonstrated significant task-induced PAC enhancement during maintenance in these three regions (AMY: p = 5.0274e-05; aHPC: p = 4.6223e-06; pHPC: p = 3.1123e-05).

For the interregional theta-gamma PAC analyses, a 2 (performance: incorrect vs. correct) × 2 (stage: baseline vs. maintenance) repeated-measures ANOVA revealed significant interaction effects for both AMY-aHPC/pHPC PAC (AMY-aHPC: p = 0.0018, F(1, 55) = 10.6819; AMY-pHPC: p = 0.0129, F(1, 50) = 6.6488) and aHPC/pHPC-AMY PAC (aHPC-AMY: p = 0.0001, F(1, 53) = 16.7314; pHPC-AMY: p = 0.0001, F(1, 49) = 17.2807; Fig. 7d), indicating that the difference in the interregional PAC between baseline and maintenance stages varied depending on whether the trials were correct or incorrect. We then performed two-tailed Wilcoxon signed-rank test to compare the interregional PAC values between baseline and maintenance. For the correct trials, the PAC values were significantly greater during maintenance for both AMY-aHPC/pHPC (AMY-aHPC: p = 2.621e-06, AMY-pHPC: p = 5.4257e-04) and aHPC/pHPC-AMY (aHPC-AMY: p = 6.6129e-07, pHPC-AMY: p = 4.1707e-05; Fig. 7d). However, for the incorrect trials, the AMY-aHPC/pHPC PAC and pHPC-AMY PAC revealed no significant differences between maintenance and baseline (AMY-aHPC: p = 0.7197, AMY-pHPC: p = 0.8882, pHPC-AMY: p = 0.6055); the aHPC-AMY showed a significant decrease PAC for the incorrect trials (p = 0.0161). Together, these results indicate that interregional theta-gamma PAC between the AMY and HPC during maintenance facilitate successful WM behavior.

## 3. Discussion

Both the AMY and HPC are involved in WM processing. Although the human HPC has been proposed to support WM maintenance in a load-dependent manner, it has remained unclear how the AMY and AMY-HPC interactions are involved in multi-item WM maintenance. In this study, we provide converging evidence that the AMY participates in multi-item WM maintenance, with their specific oscillatory patterns and interaction strengths with the HPC being load dependent. Specifically, we found that a high load elicited increased low-frequency power and intraregional theta-gamma PAC in the AMY. At the circuit level, the modulation strength of the theta phase of the HPC (including both the aHPC and pHPC) entraining the gamma amplitude of the AMY was enhanced under the high-load condition. Additionally, there was a noticeable increase in AMY-aHPC theta phase synchrony and directional connectivity strength under the high-load condition, which was directed from the aHPC to the AMY. In contrast, pHPC-AMY theta phase synchrony was not affected by memory load.

Theta rhythm has been proposed to be responsible for organizing WM items, and alpha oscillations thought to protect WM maintenance against anticipated distractors have also been associated with selective attention.^33–35^ We observed that the AMY exhibited task-induced and load-dependent low-frequency power during maintenance. Previous studies have suggested that the AMY is involved in selective attention to goal-relevant stimuli without any emotional valence.^36,37^ The increase in AMY low-frequency power during non-emotional WM maintenance, regardless of behavioral outcomes, suggests that the AMY mediates the relevance appraisal function for task-based needs.^38,39^

Within the human HPC, PAC has been recognized as a well-established mechanism for sequential memory organization and multi-item WM maintenance.^9,40^ Previous fear-learning research has demonstrated an increase in PAC in the AMY during periods of fear compared to the safe state.^41^ Our findings suggested that the phase coding mode in the AMY may not only assume a crucial role in emotional functions but also serve as a more general mechanism to support the flexible organization of complex mnemonic information.^42^

Theta synchrony would enhance information transfer and processing between AMY and HPC.^43^ Our study demonstrates a load-dependent theta phase synchrony between the AMY and HPC in correctly performed trials. A possible explanation could be that the information that is highly processed in the HPC could then be sent to the AMY, which in turn modulates neural activity in several target areas to guide attention and exert a modulatory influence on HPC-dependent memory systems.^44,45^ One popular view is that selective attention is one of the potential mechanisms involved in maintenance and influences WM performance.^46,47^ The AMY is necessary for selective attention because it serves as a central perceptual node where multimodal information converges, and the output of the AMY guides perception and covert attention and overt reorientation.^48^ In this regard, we assume that the AMY is driven to facilitate attention to task-relevant information during maintenance and that AMY-HPC interactions play an important role in incorporating WM items into neural representations. This result suggests that theta rhythm synchrony between the AMY and HPC may serve as a general integrative mechanism of cognitive circuits with divergent connectivity patterns in response to varying cognitive demands and is not limited to emotional memory consolidation.

Furthermore, our results showed that theta band information flow from the aHPC to the AMY was load dependent. It has been suggested that the HPC serves as a source of polymodal sensory information to the AMY, which has been demonstrated to act as a relevance detector in relation to current goals and needs.^49^ Some researchers considered the processes involved in WM maintenance to be primarily attentional processes.^50,51^ Considering that a high load may be related to high attentional control demands, it is plausible that the increased effective connectivity of the AMY driven by the aHPC may enhance attention to relevant stimuli and the formation of HPC-formed memories.^52^ Additionally, enhanced WM load can lead to an increase in psychological stress that has a significant impact on the activity of the AMY, primarily through the release of stress hormones. Previous human studies have shown that AMY-HPC interactions are associated with enhanced memory performance under stress.^53^ Under stress conditions, the stress hormones induced by activation of the rapid autonomic nervous system, such as noradrenaline, could enhance memory not only by directly impacting the HPC but also by indirectly modulating the HPC via the AMY.^54,55^ This modulation from the AMY enhances vigilance and increases HPC excitability, which may further improve WM maintenance under stressful conditions.

The interregional PAC is thought to serve as a mechanism for organizing neural activity across different brain regions and has been proposed to modulate synaptic plasticity.^56^ Numerous human emotional memory studies have reported PAC within the AMY and HPC. For instance, the recognition of emotional stimuli and the formation of aversive memories were associated with AMY-HPC PAC between theta and gamma oscillations,^31,57^ suggesting that information transformation via PAC between the AMY and HPC modulates multiple aspects of cognition. We provide evidence that interregional communication between the AMY and HPC via theta‒gamma PAC supports successful non-emotional WM maintenance. These results support the view that interregional PAC is a crucial neurophysiological mechanism underlying memory processing.

The interconnections between the AMY and distinct HPC regions are robust and complex. Here, our results indicated that theta phase synchrony between the AMY and HPC subregions exhibited distinct connectivity patterns. One possible explanation for the observed functional difference is that the transmembrane currents giving rise to AMY theta oscillations are likely generated by synaptic inputs from the HPC, mostly glutamatergic inputs.^58^ Classical horseradish peroxidase studies suggest that there are differences in the efferent fibers between the subregions of the HPC, with the ventral HPC preferentially projecting to the AMY.^24^ Considering the essential role of the HPC in the attention process,^59^ particularly the key role of afferent projections from the ventral HPC in the cognitive process of selective attention in rodents,^60^ a plausible explanation for the aHPC-to-AMY information flow might be its involvement in the control of selective attention under the high-load condition.

While observing the load-dependent theta phase synchrony only between the aHPC and AMY, we also found significant and intense modulation of interregional theta-gamma PAC for both the aHPC-AMY and pHPC-AMY under the high-load condition. There is still no consensus on whether these two phenomena are directly related to each other. One previous study suggested that both same-frequency coupling and cross-frequency coupling may be part of the same mechanism for interregional communication.^61^ In contrast, other studies have reported no strong overlap between phase synchrony and interregional PAC.^62^ Our results suggest that the interregional PAC may not be directly explained by theta phase synchrony, supporting the opinion that it is necessary to evaluate multiple coupling measures to quantify interregional correlation.^63^

These findings indicate a notable engagement of the AMY during non-emotional WM maintenance, showing a low-frequency information flow from the aHPC to the AMY and an enhanced strength of HPC theta phase entraining AMY gamma amplitude during high load condition. This observation contrasts with the established literature, which predominantly emphasizes the role of the AMY in emotional processing, wherein information typically flows from the AMY to the HPC ^57^, accompanied by AMY-HPC theta-gamma coupling^31,64^. This suggests a potential framework in which the AMY may not only be a center for emotional responses but also actively participates in organizing and maintaining information under varying cognitive demands. Within this framework, the HPC provides processed polymodal sensory information to the AMY, enabling it to modulate neural activity across various target areas, thereby guiding attention and influencing HPC-dependent memory systems. Given the increased cognitive demands associated with high WM load, the effective connectivity between the AMY and HPC enhances attentional control and promotes the maintenance of relevant information. Future research should explore the specific neural interactions that enable the AMY to facilitate non-emotional WM and examine its potential implications for understanding the complex mechanisms where emotional and cognitive processes converge. This may provide valuable insights for developing treatment strategies for memory-related disorders.

In conclusion, our study offers evidence supporting the functional involvement of the AMY in non-emotional higher-level cognitive processes and provides novel insight into the neurophysiological basis of the human AMY and AMY-HPC interactions during WM maintenance.

## 4. Methods

### 4.1. Subjects and electrodes

We analyzed an iEEG dataset from nine right-handed patients, all of whom were diagnosed with drug-resistant focal epilepsy.^25^ These patients had intracranial depth electrodes (1.3 mm diameter, 8 contacts of 1.6 mm length, spacing between contact centers 5 mm, ADTech®, Racine, WI, www.adtechmedical.com) implanted stereotactically for potential surgical treatment of epilepsy. The configurations for electrode implantation were determined solely based on clinical needs. Each subject provided informed consent before the experiment. This study was approved by the institutional ethics review board (Kantonale Ethikkommission Zürich, PB-2016-02055).

### 4.2. Data acquisition and electrode localization

The iEEG recordings were performed against a common intracranial reference via the Neuralynx ATLAS recording system (0.5-5000 Hz passband, Neuralynx®, Bozeman, MT, USA). The sampling rate of the iEEG recordings was 2000 Hz. Electrodes were localized in each subject using post-implantation computed tomography (CT) scans and post-implantation structural T1-weighted magnetic resonance imaging (MRI) scans. The dataset included the normalized Montreal Neurological Institute (MNI) coordinates of each contact and corresponding anatomical labels assigned by the Brainnetome Atlas.^65^

### 4.3 Channel selection and pre-processing

Due to clinical considerations, the targeted regions and hemispheres varied among the subjects (Table 1). To minimize interindividual variability, the two most medial channels were selected on each electrode targeting the AMY, aHPC and pHPC in each hemisphere, in accordance with previous studies.^66^ In total, we identified 32 aHPC, 32 pHPC and 30 AMY contacts across all subjects. For the interregional analysis, only ipsilateral channel pairs were included. Across all subjects, the number of electrode pairs covering the AMY-aHPC within the same hemisphere was 60, and the number of electrode pairs covering the AMY-pHPC was 56.

**Table 1.** Number of intracranial contacts for the AMY, aHPC and pHPC.

| Subject | AMY (left) | AMY (right) | aHPC (left) | aHPC (right) | pHPC (left) | pHPC (right) |
| --- | --- | --- | --- | --- | --- | --- |
| 1 | 8 | 0 | 8 | 0 | 8 | 8 |
| 2 | 8 | 8 | 8 | 8 | 8 | 8 |
| 3 | 8 | 0 | 8 | 8 | 8 | 0 |
| 4 | 8 | 8 | 8 | 8 | 8 | 8 |
| 5 | 8 | 0 | 8 | 0 | 0 | 8 |
| 6 | 8 | 8 | 8 | 8 | 8 | 8 |
| 7 | 8 | 8 | 8 | 8 | 8 | 8 |
| 8 | 8 | 8 | 8 | 8 | 8 | 7 |
| 9 | 8 | 8 | 8 | 8 | 8 | 8 |

Data pre-processing was performed using the EEGLAB^67^ and FieldTrip (http://fieldtrip.fcdonders.nl/) toolboxes and custom MATLAB (MathWorks, Natick MA, USA) scripts. First, we performed bipolar re-referencing by subtracting the signal from the contact closest to the channel of interest. Bipolar re-referencing has been suggested to effectively improve sensitivity and reduce artifacts by eliminating distant signals picked up by adjacent electrode contacts.^68^ After bipolar re-referencing, the iEEG recordings were bandpass filtered between 1−200 Hz using a fourth-order Butterworth filter with a Hamming window. Only trials with correct behavioral responses were included in our analysis of the load effect. Then, we concatenated these trials and Z-transformed the signals of each channel separately by removing the mean and scaling by the standard deviation. We divided the continuous data into epochs based on the trials and then rejected any trials that were determined to contain artifacts by visual inspection (82.6% of all correct epochs remained in the analysis).

### 4.4. Spectral power analysis

Time-frequency representations of power were computed for low-load (set size 4) and high-load (set size 6) trials in the AMY, aHPC and pHPC. We used the wavelet method (with 6 cycles) implemented in FieldTrip. We computed the power in 1 Hz steps across frequencies ranging from 3 Hz to 100 Hz with a temporal resolution of 0.01 s. The power from 0.5 to 0 s before stimulus onset served as the baseline power. We corrected the baseline by transforming the power estimates to decibels relative to the baseline power. When comparing the normalized power at each time-frequency point between different load conditions, we used cluster-based nonparametric permutation tests (with 2000 permutations). Then, theta and alpha power were calculated by averaging the normalized power over 3–8 Hz and 8–13 Hz, respectively. A Wilcoxon signed-rank test (two-tailed) was used to compare the low-frequency power between the two load conditions.

### 4.5. Interregional phase synchrony

To evaluate the functional connectivity between the AMY and the aHPC/pHPC, we calculated the wPLI, which is less sensitive to volume conduction driven by a single or shared source.^29^ The wPLI is based solely on the imaginary part of the cross-spectrum between two signals and has been shown to exhibit the best performance in the presence of noise compared to other phase statistics.^69^ The wPLI was computed as follows:

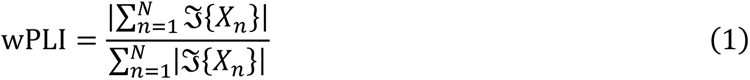

where N is the total number of trials and ℑ{𝑋_n_} represents the imaginary part of the cross-spectrum in the nth trial. We used the same time-frequency decomposition parameters as those used for the spectral power analysis to obtain phase information. The wPLI ranges from 0 to 1, with values approaching 1 indicating stronger phase synchrony.

To avoid any bias due to the different number of trials for the two load conditions, the trial number was matched to the lower number of the two conditions by random selection. Before calculating the wPLI for each electrode pair and condition, the trial-wise mean was subtracted from each trial to minimize contamination from simultaneous voltage changes on phase estimations. To test the statistical significance of the wPLI, we shuffled trials of all conditions 200 times to create a null distribution and computed the 95th percentile threshold. We averaged the null distribution of all electrode pairs and compared the observed wPLI with this null distribution. Only electrode pairs with wPLI values above the corresponding threshold were selected for further analysis. Next, we extracted and averaged the wPLI within the theta (3–8 Hz) and alpha (8–13 Hz) bands and compared the wPLI estimates for the low- and high-load conditions within each band via the paired Wilcoxon signed-rank test (two-tailed).

### 4.6. Granger causality analysis

To examine the directionality of the AMY-aHPC and AMY-pHPC synchrony, we analyzed the effective connectivity in the theta band during the maintenance period. We used the Multivariate Granger Causality (MVGC) Toolbox (based on the Akaike information criterion to define the model order) to compute the spectral GC within the theta band (3–8 Hz) for the electrode pairs with significant theta phase synchrony.^70^ The GC index was computed for the low- and high-load trials separately. Significance testing was conducted by creating the null distribution of GC by randomly shuffling and switching directions across trials for each electrode pair 200 times. The GC estimates were considered significant if they exceeded the 95th percentile of the null distribution. We extracted the GC index from significant electrode pairs for both directions within each circuit (AMY-aHPC: AMY to aHPC, aHPC to AMY; AMY-pHPC: AMY to pHPC, pHPC to AMY). To estimate the effect of load on the effective connectivity strength, we used the Wilcoxon signed-rank test (two-tailed) to compare the GC index between low- and high-load trials for each direction of each circuit.

Within-frequency phase-slope index To reduce spurious connectivity arising from volume conduction effects and provide a more robust measurement, we re-assessed the interregional directionality in the theta band using the PSI from both directions for the aHPC-AMY and pHPC-AMY circuits. First, the trial-wise mean was subtracted from each trial of the low- and high-load conditions. Then, the PSI was computed for each electrode pair from different brain regions in the frequency band of 3–8 Hz during the WM maintenance period as described by Nolte et al.^30^ The PSI estimates the slope of the phase difference between the sender signals and receiver signals; a positive PSI indicates that the seed region acts as a sender (modulating signals), and a negative PSI indicates that the seed region acts as a receiver (modulated signals). To assess the significance level of the PSI, we randomly shuffled the trials 200 times to generate a null distribution for each electrode pair. The raw PSIs were z-scored on the null distribution. A significant PSI was defined as |z| > 1.96 (i.e., α = 0.05). Then, we extracted the electrode pairs with significant z-scored PSIs for further analysis. To confirm whether the directional information flow was modulated by WM load, we compared the PSI values between the low- and high-load conditions using the Wilcoxon signed-rank test (two-tailed).

Intra- and interregional cross-frequency phase amplitude coupling analysis The cross-frequency PAC was estimated within and between regions using Tort’s modulation index (MI) method.^71^ We estimated the phase of low frequencies between 2 and 10 Hz (in steps of 1 Hz; termed “low-frequency phase”) and the amplitude of frequencies between 30 and 100 Hz (in steps of 5 Hz; termed “high-frequency amplitude”) and computed the MI under both conditions. In brief, we obtained the instantaneous power and phase representations using the Morlet convolution with a width of seven cycles for each epoch. Then, the maintenance intervals were extracted and concatenated into one continual EEG signal for each channel. The low-frequency phases were divided into 20 bins (i.e., 18° per phase bin), and the mean amplitude of high-frequency oscillations in each bin was calculated. The distribution across bins was uniform if the PAC was not evident. A normalized version of the Kullback‒Leibler divergence was used to measure the distribution of this histogram and determine MI values. This procedure was repeated for all phase–amplitude frequency pairs. For interregional PAC, we performed two directions estimates—the theta phase of the AMY modulating the gamma amplitude of the aHPC/pHPC and the theta phase of the aHPC/pHPC modulating the gamma amplitude of the AMY—for the low- and high-load conditions.

To test the significance of the raw PAC, we used a surrogate test, as in previous studies. Specifically, we randomly generated the time lag of the raw phase time series and the temporal relationship between the phase and amplitude information and recomputed the PAC using the shuffled phase time series. This procedure was repeated 200 times to create the null distribution of PAC values. The raw PAC values were then z-scored against the null distribution to correct for any spurious results and evaluate the significance of the PAC values. The z-scored PAC values that exceeded a threshold value of 1.96 were considered significant.^62^ Only single channels from target regions (for intraregional PAC estimates) or electrode pairs from two regions within the same hemisphere (for interregional PAC estimates) exhibiting significant PAC values were included in subsequent analyses. We then averaged the theta (3–8 Hz)-gamma (30–100 Hz) z-scored PAC values for the low- and high-load trials. A Wilcoxon signed-rank test (two-tailed) was applied to estimate the effect of load on the intraregional PAC and interregional PAC.

## Data availability

The raw data used in this study have been downloaded from a public database^25^ at the following link: https://doi.gin.g-node.org/10.12751/g-node.d76994/.

